# Electrophysiological Frequency Band Ratio Measures Conflate Periodic and Aperiodic Neural Activity

**DOI:** 10.1101/2020.01.11.900977

**Authors:** Thomas Donoghue, Julio Dominguez, Bradley Voytek

**Affiliations:** Department of Cognitive Science, University of California, San Diego; Halıcıoğlu Data Science Institute, University of California, San Diego; Neurosciences Graduate Program, University of California, San Diego

**Keywords:** neural oscillations, frequency band ratios, spectral power ratios, theta / beta ratio, theta / alpha ratio, alpha / beta ratio, electroencephalography, 1/f activity, aperiodic neural activity

## Abstract

A common analysis measure for neuro-electrophysiological recordings is to compute the power ratio between two frequency bands. Applications of band ratio measures include investigations of cognitive processes as well as biomarkers for conditions such as attention-deficit hyperactivity disorder. Band ratio measures are typically interpreted as reflecting quantitative measures of periodic, or oscillatory, activity, which implicitly assumes that a ratio is measuring the relative powers of two distinct periodic components that are well captured by predefined frequency ranges. However, electrophysiological signals contain periodic components and a 1/f-like aperiodic component, which contributes power across all frequencies. In this work, we investigate whether band ratio measures reflect power differences between two oscillations, as intended. We examine to what extent ratios may instead reflect other periodic changes—such as in center frequency or bandwidth—and/or aperiodic activity. We test this first in simulation, exploring how band ratio measures relate to changes in multiple spectral features. In simulation, we show how multiple periodic and aperiodic features affect band ratio measures. We then validate these findings in a large electroencephalography (EEG) dataset, comparing band ratio measures to parameterizations of power spectral features. In EEG, we find that multiple disparate features influence ratio measures. For example, the commonly applied theta / beta ratio is most reflective of differences in aperiodic activity, and not oscillatory theta or beta power. Collectively, we show how periodic and aperiodic features can drive the same observed changes in band ratio measures. Our results demonstrate how ratio measures reflect different features in different contexts, inconsistent with their typical interpretations. We conclude that band ratio measures are non-specific, conflating multiple possible underlying spectral changes. Explicit parameterization of neural power spectra is better able to provide measurement specificity, elucidating which components of the data change in what ways, allowing for more appropriate physiological interpretations.

**Materials Descriptions & Availability Statements:** *Project Repository:* This project is also made openly available through an online project repository in which the code and data are made available, with step-by-step guides through the analyses. Project Repository: http://github.com/voytekresearch/BandRatios

*Datasets:* This project uses simulated data, literature text mining data, and electroencephalography data. Simulated Data
The simulations used in this project are created with openly available software packages. Settings and code to re-generate simulated data is available with the open-access code for the project. Copies of the simulated data that were used in this investigation are available in the project repository. Literature Data
Literature data for this project was collected from the PubMed database. Exact search terms used to collect the data are available in the project repository. The exact data collected from the literature and meta-data about the collection are saved and available in the project repository. EEG Data
The EEG data used in this project is from the openly available dataset, the ‘Multimodal Resource for Studying Information processing in the Developing Brain’ (MIPDB) database. This dataset is created and released by the Childmind Institute. This dataset was released and is re-used here under the terms of the Creative Commons-Attribution-Non-Commercial-Share-Alike License (CC-BY-NC-SA), and is described in (Langer et al., 2017).
Child Mind Institute: https://childmind.org
Data Portal: http://fcon_1000.projects.nitrc.org/indi/cmi_eeg/

*Software:* Code used and written for this project was written in the Python programming language. All the code used within this project is deposited in the project repository and is made openly available and licensed for re-use. As well as standard library Python, this project uses 3^rd^ party software packages *numpy* and *pandas* for data management, *scipy* for data processing, *matplotlib* and *seaborn* for data visualization and *MNE* for managing and pre-processing data. This project also uses open-source Python packages developed and released by the authors: Simulations and spectral parameterization were done using the FOOOF toolbox. Code Repository: https://github.com/fooof-tools/fooof Literature collection and analyses were done using the LISC toolbox. Code Repository: https://github.com/lisc-tools/lisc

## Introduction

### 1.1 History & Introduction of Band Ratio Measures

Studies in cognitive and clinical neuroscience employ a broad range of analyses that are designed to measure how electrophysiological measures vary with, and potentially predict, features of interest such as behavioral outputs and disease states. Many such analyses focus on putative rhythmic, or oscillatory, activity, organized into distinct frequency bands such as theta, alpha and beta, that will collectively be referred to as ‘periodic’ activity. One such analysis method is to calculate the ratio of power between two of these pre-specified frequency bands. For example, the theta / beta ratio is calculated as the average power in the theta band, typically 4-8 Hz, divided by the average power in the beta band, typically within the range of 13-30 Hz. Such measures can be applied to electroencephalography (EEG), magnetoencephalography (MEG), electrocorticography (ECoG) and/or local field potential (LFP) data and have been argued to be a biomarker for a variety of cognitive correlates (for example, attentional control: Angelidis, van der Does, Schakel, & Putman, 2016), and clinical disorders (for example, ADHD: Arns, Conners, & Kraemer, 2013; or Alzheimer’s: Cassani, Estarellas, San-Martin, Fraga, & Falk, 2018).

An early example of such an approach was to measure, from the correllelogram of EEG data, the ratio of the dominant rhythm to the ‘background’ activity (Daniel, 1964). This measure was developed to leverage emerging tools for spectral analysis to quantify electrophysiological features of interest and integrate computational approaches, in what would later come to be referred to as ‘quantitative EEG’ or ‘qEEG’. As spectral power estimation procedures became more common, studies began using frequency band ratios calculated directly from estimations of band powers extracted from power spectra, such as the ratio of theta to alpha power (Matoušek, 1968), which is now the standard approach for calculating frequency band ratio measures (see Figure 1A).

**Figure 1.**
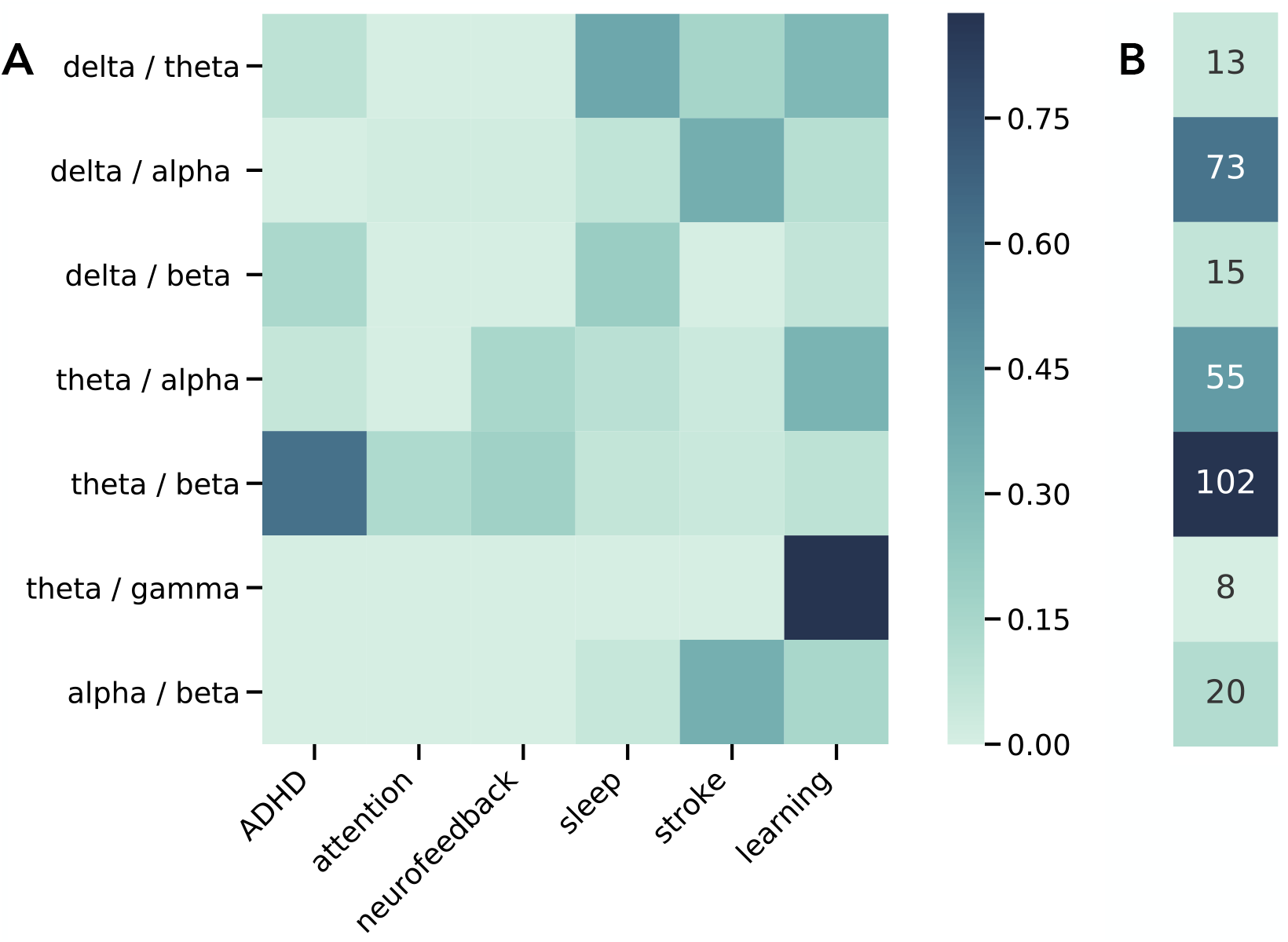
Literature Analysis of Band Ratio Related Articles. **A)** Associations between published journal articles referring to band ratio measures and cognitive and clinical associations. Each cell represents the proportion of articles referring to a specified band ratio measure that also mentions the corresponding association term. **B)** Total counts of the number of articles mentioning each band ratio measure.

Early work used band ratio measures because they were found to be more stable than either absolute or relative measures of individual frequency band powers (Daniel, 1964; Matoušek, 1968). Relative power measures, including band ratios, are also used as a data normalization method, to control for potential differences in confounds such as skull thickness and volume conduction, that otherwise make absolute measures difficult to compare and interpret across individuals. Several investigations also reported correlated changes between frequency bands, such as a frequency ‘slowing’, whereby low frequency power increases and high frequency power decreases, and therefore recommended frequency band ratio measures as an ideal measure to capture such changes (Lubar, 1991).

### 1.2 Applications of Band Ratio Measures

In cognitive neuroscience, band ratio measures are often used in EEG studies investigating possible physiological correlates of behaviors of interest, including investigations exploring vigilance and alertness (Matoušek & Petersén, 1983), cognitive development and aging (Clarke et al., 2001), reward processing (Schutter & Van Honk, 2005), and affect (Putman et al., 2010). One of the most consistent lines of research in this area focuses on the theta / beta ratio as a potential biomarker for executive function, and in particular attentional processing (Angelidis et al., 2016; Gordon et al., 2018; Lubar, 1991), with recent reports investigating, for example, cognitive control (Angelidis et al., 2018), and attentional control (van Son et al., 2019). Other work using EEG experiments have explored ratio measures in learning and memory, examining, for example, short term memory using the theta / beta ratio (Trammell et al., 2017), and memory impairment using the theta / gamma ratio (Moretti et al., 2009). Similar work in animals has investigated the theta / delta ratio in hippocampal recordings during associative learning paradigms in rabbits (Nokia et al., 2008) and rats (Kim et al., 2016).

Frequency band ratio measures have also been used to explore changes within and between individuals in contexts such as state mapping and sleep scoring, and work in development and aging. In developmental work, ratio measures have been included in investigations of age related electrophysiological changes (Clarke et al., 2001; Gasser et al., 1988; Matoušek & Petersén, 1973). Several proposed approaches for automated sleep stage classification have also used band ratio measures and found them to be useful measures (Costa-Miserachs et al., 2003; Krakovská & Mezeiová, 2011; Reed et al., 2017; van Luijtelaar & Coenen, 1984). This includes work using the theta / delta ratio for sleep scoring of hippocampal local field data in rats (Costa-Miserachs et al., 2003; van Luijtelaar & Coenen, 1984), and delta / beta ratio for human data analysis, including EEG (Krakovská & Mezeiová, 2011) and ECoG (Reed et al., 2017).

In clinical neuroscience, band ratios are also a common approach, including in studies seeking biomarkers for diagnosis, clinical monitoring, and potential intervention. Investigations into the potential clinical utility of band ratio measure include investigations of anesthesia (Long et al., 1989), disorders of consciousness (Pfurtscheller et al., 1986), multiple sclerosis (Keune et al., 2017), cerebral ischemia (Sheorajpanday et al., 2009), and Parkinson’s disease (Geraedts et al., 2018). In psychiatry, band ratios measures have been applied in studies of autism (Wang et al., 2016) and as a potential biomarker for psychotic disorders (Howells et al., 2018). Band ratios are also commonly investigated in the search for biomarkers for mild-cognitive impairment, dementia, and Alzheimer’s (Bennys et al., 2001; Moretti et al., 2013; Penttilä et al., 1985), recently reviewed in (Cassani et al., 2018).

The most common clinical application of band ratios measures is in investigations of attention-deficit hyperactivity disorder (ADHD) (Loo & Makeig, 2012). After early work reported a relative increase in theta and decrease in beta in ADHD, theta / beta ratios were proposed as a potential biomarker for the disorder (Lubar, 1991), which prompted a large number of studies investigating the theta / beta ratio as a descriptive feature and potential diagnostic biomarker for ADHD (see reviews in Arns, Conners, & Kraemer, 2013 & Snyder & Hall, 2006). Initial work was very promising with an early meta-analysis supporting that theta / beta ratios may be a predictive marker of ADHD, reporting a pooled effect size of 3.08 (Snyder & Hall, 2006). However, a more recent review found much weaker support for this conclusion, also reporting that the effect size of theta / beta ratios differentiating between ADHD and control groups has been decreasing across time (Arns et al., 2013). Notably, the theta / beta ratio has been investigated as a potential diagnostic marker of ADHD (Snyder et al., 2015), and although this led to approval by the Food and Drug Administration of the United States, inconsistent evidence regarding the efficacy of using the theta / beta ratio in diagnostic practice has led to a practice advisory against using it (Gloss et al., 2016).

As well as being used in investigations seeking diagnostic biomarkers, band ratio measures are commonly targeted in neurofeedback paradigms. This includes clinical applications using protocols aimed at manipulating theta / beta ratio for the treatment of ADHD (Arns et al., 2014), and as potential treatments for disorders such as autism (Wang et al., 2016). Non-clinically related neurofeedback protocols using band ratio measures have also been explored, including investigations aimed at manipulating and improving attentional and executive functions (Studer et al., 2014; Vernon et al., 2003) and relaxation (Egner et al., 2002; Raymond et al., 2005).

Collectively, band ratio measures are used across basic, clinical, and applied neuroscience to examine a wide variety of their correlates. To explore the breadth of reported band ratio correlations, we also ran an automated literature search that collects information on the number of published articles that reference each ratio term and their major associates (Figure 1). This analysis shows that theta / beta ratio measures are the most common, though a variety of other band ratios are commonly applied, with distinct applications. We find over 250 articles that mention band ratio measures, supporting that these are a relatively common method to apply to electrophysiological data, across a wide range of applications. This is also likely an underestimate, as our text-mining approach is limited to specific phrases that appear only in article abstracts.

### 1.3 Methodological Properties & Interpretations of Band Ratio Measures

Given the popularity of band ratios across domains, and their reported clinical utility, it is important to investigate and understand the properties and assumptions of such analyses, and how those assumptions relate to their interpretations. In this investigation we examine whether the general conception of band ratios as measures that specifically reflect periodic neural activity is well founded in the face of work showing that periodic properties of electrophysiological data are highly variable, often violating the assumptions of predefined frequency bands, and that they co-exist with variable and dynamic aperiodic activity (Haller et al., 2018).

Methodologically, studies using band ratios typically follow a stereotyped procedure whereby power in pre-defined, fixed frequency bands are calculated, from which a ratio is calculated. Band ratios are typically calculated from absolute power values, though some studies use relative or normalized power measures in which the power within a band is normalized by total power. Because ratios typically display a non-normal, skewed distribution, they are often log-transformed before further analysis.

This ratio measure is then used as an electrophysiological marker that is then either analyzed for potential correlations with features of interest, and/or used as a target in neurofeedback paradigms. Band ratio measures are often conceptualized as capturing the proportion of a ‘slower’ frequency band relative to some ‘faster’ one, and are often interpreted as a relative ‘slowing’ of neural activity (eg: Monastra, Lubar, & Linden, 2001; Poza, Hornero, Abásolo, Fernández, & Mayo, 2008) or as a shift of power from one band to another (eg: Gasser, Verleger, Bächer, & Sroka, 1988), which conceptualize one process explaining a change in periodic activity. Other interpretations focus on interpreting and investigating ratio measures more in terms of changes within the component bands, for example interpreting a decrease in theta / beta ratio as changes in the theta or beta band (eg: Clarke et al., 2013), which conceptualizes one or more distinct changes in periodic bands. Differences in these such interpretations include whether ratio measures are conceptualized to reflect one change of reapportioning activity, or potentially multiple changes in distinct bands.

What is common across these conceptualizations is that they interpret ratio measures as reflecting periodic power, and so presume, as many investigations do, that pre-specified frequency bands specifically measure periodic oscillatory activity. For this assumption to be valid, defined frequency bands of interest, for example, 4-8 Hz theta, must capture periodic activity that is to be considered as relating to that band. A known problem with applying predefined frequency bands uniformly across all participants is that variation in center frequencies can lead to misestimations of the desired features. This can be an underestimation, if frequency variation causes band power to ‘move’ outside the canonical range, or an overestimation, if power from an adjacent frequency band is captured in the examined range. Similar issues can arise if the bandwidth of frequency bands violates expectations and/or is different between groups. These potential periodic confounds challenge the assumption that band ratio measures relate specifically to relative periodic power (see Figure 3A).

An example of this issue has been previously demonstrated in a sample of participants with ADHD, whereby an increased theta / beta ratio, as measured using canonical band definitions, was found to actually reflect a slowed alpha peak in the ADHD group (Lansbergen et al., 2011). In this case, the theta / beta ratio calculated using individualized frequency bands found no difference between groups. This suggests that, in at least some cases, frequency variation can lead to measurements and interpretations of band ratios that do not accurately reflect the actual properties of the data. This has led to suggestions that band ratios measure should be computed using individualized frequency bands (Saad et al., 2018).

Beyond the periodic confounds, a broader issue is the implicit assumption that frequency definitions reflect periodic activity in the data, and that this activity can be specifically captured by measuring power averaged across a frequency range. This assumption is in general invalid, as electrophysiological activity includes not only periodic components, but a 1/f-like distributed aperiodic component (Haller et al., 2018; B. J. He, 2014), which has power at all frequencies, but does not consist of periodic activity (see Figure 2B). The presence of this 1/f-like activity, henceforth referred to as the ‘aperiodic component’, entails that there will always be power in a given frequency range, but that this power should not necessarily be assumed to reflect periodic activity. Rather, power at a particular frequency, or frequency range, reflects, at least in part, aperiodic activity, and only partially, if at all, reflects periodic activity. A marker that there is actual periodic power in a signal is that there should be a band specific peak over and above this aperiodic component (Buzsáki et al., 2013). To specifically measure this periodic component of the signal, one should measure the power in this band specific peak relative to the aperiodic component of the signal (Haller et al., 2018).

**Figure 2.**
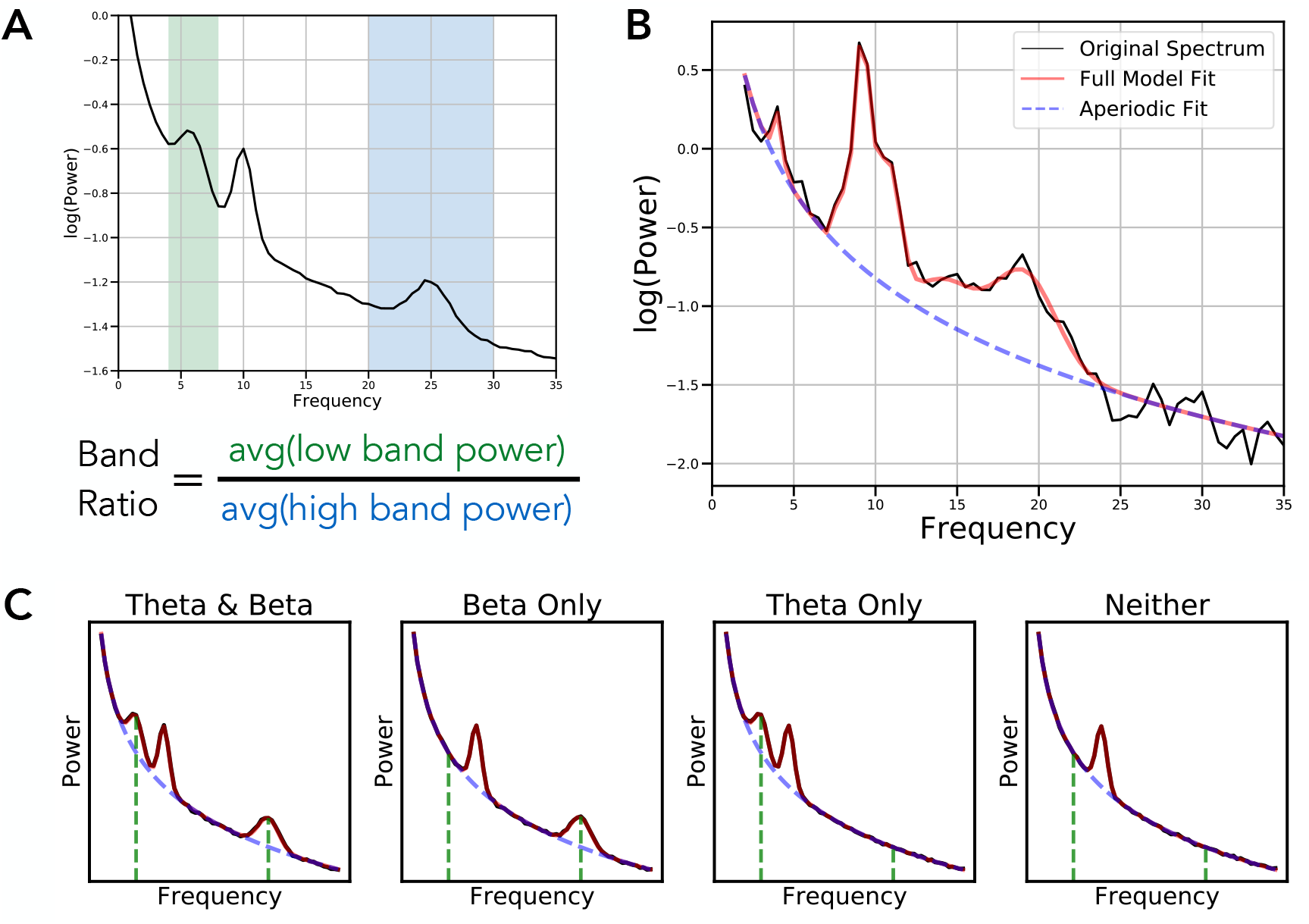
Overview of Band Ratio Measures and Spectral Parameters. **A)** An example power spectrum in which shaded regions reflect the theta (4-8 Hz) and beta band (20-30 Hz) respectively. Band ratio measures, such as the theta / beta ratio are taken by dividing the average power between these two bands. **B)** An example of a parameterized power spectrum, in which aperiodic activity is separated from measured periodic components. **C)** Examples of simulated power spectra with and without component oscillations of the theta / beta ratio. Black lines indicate the simulated data, with red line reflecting the model fit, the dashed blue line indicating the aperiodic component of the model fit, and the green lines indicating the location of canonical theta and beta oscillations. Band ratio measures, though intended to measure periodic activity, will reflect power at the pre-determined frequencies regardless of whether there is evidence of periodic activity at these frequencies.

**Figure 3.**
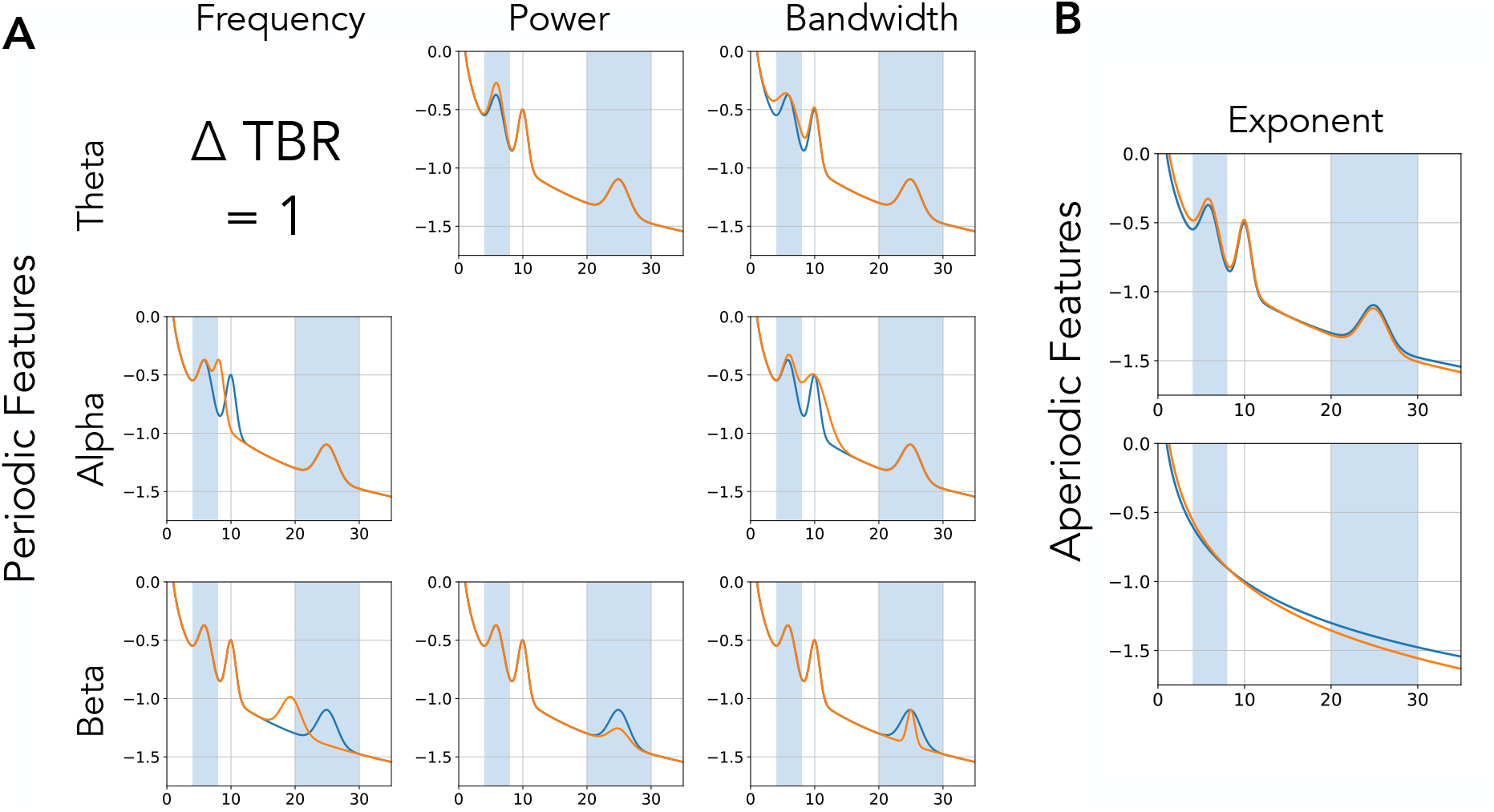
Equivalent Band Ratio Differences from Distinct Changes. Simulations demonstrating the underdetermined nature of band ratio measures. In each case, the power spectrum plotted orange has the same difference of measured theta / beta ratio from the reference spectrum, in blue. This difference in ratio can arise from changes in multiple different features of the data, including a shift in: **A)** the periodic properties such as the center frequency, power or bandwidth of oscillations, and/or from a shift in; **B)** aperiodic component of the data. Differences in aperiodic activity can induce differences in measured band ratios, even without any periodic components present.

Band ratio measures, as currently applied, do not address the confound of ubiquitous aperiodic activity in neural signals. Aperiodic neural activity is known to be variable both within (Podvalny et al., 2015) and between individuals (Voytek et al., 2015). This variability raises the possibility that band ratio measures may reflect, at least partially, aperiodic activity and that measured differences within and between individuals may be driven by differences in aperiodic properties of the data (see Figure 3B). The very observation that there are correlated changes across frequency bands that helped popularize band ratio measures (Lubar, 1991) can even be interpreted to support the suggestion that a parsimonious description of the data could be changes in aperiodic properties, across all frequencies. This is also broadly consistent with the interpretations of ratios reflecting ‘substitutions’ of power between bands (Gasser et al., 1988) in the sense that one process explains the changes across different frequency regions (though inconsistent with this being a shift of periodic activity).

In summary, band ratio measures are a common analyses measure that are calculated across two frequency bands that are designed to, and are interpreted as, reflecting relative periodic activity. However, even when oscillations are clearly present, variations in the measure may reflect not only the power across the two bands, but may be driven by differences in the center frequencies, and/or the bandwidths of such periodic components and, can also be driven by changes in aperiodic activity with or without periodic activity being present (see Figure 2C). Altogether, this suggests that band ratio measures are underdetermined, whereby a change in one or many different features of the data may drive analogous differences in band ratio measures (Figure 3). If so, not only are typically interpretations of band ratio measures unsupported, but band ratio measures, by themselves, may be essentially uninterpretable, as underlying physiological causes of changes in the measure are undecipherable from the measure itself, but reflect different properties of the data.

To investigate these issues, we examine the properties and validity of band ratio measures, including, 1) how are band ratio measures influenced by different features of periodic activity, including center frequency, power and bandwidth, and 2) how band ratio measures are influenced by changes in aperiodic properties of the data, including the aperiodic exponent and offset. We start by systematically exploring the properties of band ratio measures across simulated data that mimic the statistics of real data, for which ground truth is known. We use these simulations to evaluate how changes in different features, and their combinations, influence band ratio measures. We follow by analyzing a large EEG dataset (*n* = 126) in which we applied band ratio measures and compared ratios to methods that explicitly parameterize periodic and aperiodic features of the data, to infer which neural features influence and contribute to band ratio measures. We find that many different features of the data can give rise to band ratio differences, making them effectively uninterpretable in isolation, without additional context of the rest of the power spectral features involved. Therefore, we conclude that band ratios should not be interpreted as a well-posed method to specifically measure periodic properties of neural times series, and comment on how the methodological findings from this work can be used to interpret prior work, and what it suggests for future investigations.

## Methods

In order to investigate the properties of frequency band ratios, we explored calculating band ratio measures across simulated power spectra, for which ground truth values were known, as well as investigating their application in EEG data. As a comparison to the band ratio measures, periodic (oscillatory) and aperiodic properties of power spectra were characterized using the fitting oscillations-&-one-over-f (FOOOF) toolbox (Haller et al., 2018). Band ratio measures were compared to the outputs of the parameterization of the power spectra, which quantifies the center frequency (CF), power (PW) and bandwidth (BW) of identified periodic components, as well as the exponent and offset (described below) of the aperiodic component. Using these parameterizations, we evaluate which components of the data the band ratio measures reflect. For all analyses, canonical frequency band definitions were defined as: theta (4-8 Hz), alpha (8-13 Hz), beta (13-30 Hz).

Analyses were done using Python (version 3.7), including common libraries numpy, pandas, scipy, matplotlib and seaborn for analysis and visualization. The MNE library was used for managing and processing EEG data (Gramfort et al., 2014). Custom code was used to calculate band ratio measures and perform analyses. All code for this project is available in the project repository (https://github.com/voytekresearch/BandRatios).

### 2.1 Literature Analysis

The literature analysis was done use the ‘Literature Scanner’ (LISC) Python toolbox (Donoghue, 2019). Briefly, this toolbox allows for collecting and analyzing literature data by curating search terms of interest, gathering related articles from available databases, and analyzing the results. For this analysis, a list of band ratio terms (e.g., “theta / beta ratio”) and related association terms (e.g., “attention”), with relevant synonyms and exclusion words, was manually curated. Searches were performed to determine the number of articles in the PubMed database that reference these terms in their abstract, and the number of co-occurrences of band ratio terms with association terms. Association scores were calculated as the proportion of articles referencing a band ratio measure that also mention one of the included association terms.

### 2.2 Simulations

Neural power spectra were simulated to match the statistics of electrophysiological neural data, by combining a 1/f-like aperiodic component with overlying peaks of periodic activity, with overlying noise (Haller et al., 2018). The aperiodic component describes the 1/f-like characteristic of neural power spectra and is entirely described by the aperiodic ‘exponent’ and ‘offset.’ The aperiodic exponent, meaning the *χ* in 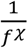, describes the steepness of the 1/f, and the ‘offset,’ describes the vertical translation of the aperiodic activity. Periodic components describe putative oscillations which display power above the aperiodic component. Periodic components are simulated as Gaussians, and are described by a ‘center frequency’ (CF) in hertz, ‘power’ (PW) from the aperiodic component to the oscillatory peak in arbitrary units (au), and ‘bandwidth’ (BW) which describes the width of the peak, also measured in hertz. The simulation ultimately follows:

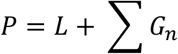

in which *L* is the aperiodic component, described as

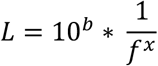

in which b is the offset and *χ* is the exponent. On top of this, periodic components are added with each of *n* peaks described as a Gaussian, as:

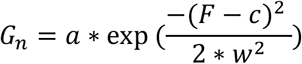

in which *c* is the peak center frequency, and *a* and *w* are the height and width of the gaussian, equivalent to the power and bandwidth of the peak.

Spectra were simulated for the frequency range of 1-35 Hz, with a 0.5 Hz frequency resolution. Default aperiodic and periodic parameter values were chosen to capture physiologically realistic values. A small amount of normally distributed noise (0.005 au) was added to all spectra to emulate real power spectra.

We calculated band ratios from simulated power spectra by dividing mean power across the low band range by the mean power across a high band range. We calculated the theta / beta ratio, theta / alpha ratio, and alpha / beta ratio.

To measure how spectral parameters relate to band ratio measures, spectra were simulated where a single parameter was varied across a range while the remaining parameters were kept at their default values. From these spectra the theta / beta, theta / alpha and alpha / beta ratios were calculated to track how individual parameters affect ratio measures. Since CF, PW, and BW are specific to a peak, they were all individually varied for both low-band and high-band peaks. The ranges of values for each parameter are given in supplemental tables 1 & 2.

We then studied how band ratio measures are affected by multiple interacting changes in spectral parameters. Further simulations were carried out as two parameters from the set {CF, PW, BW, EXP} were simultaneously varied across their respective ranges. All combinations of paired parameter simulations were calculated and analyzed. The default parameter settings and ranges remained the same as the single parameter simulations.

### 2.3 EEG Data Analysis

To study how various spectral parameters affect band ratio measures, we used the openly available ‘Multimodal Resource for Studying Information Processing in the Developing Brain’, or MIPDB, dataset of human EEG data released by the Child Mind Institute (Langer et al., 2017). The study population is a community sample of children and adults (*n* = 126, age range = 6-44, age mean = 15.79, age standard deviation = 8.03, number of males = 69). Data for each subject includes resting state and task EEG data, behavioral measures, and eye tracking data. For the current investigation, we analyzed eyes-closed resting state data, collected on a 128 channel Geodesic Hydrocel system. Of the 126 participants in the dataset, 9 did not include resting state data collection, as indicated by the dataset description, and were therefore excluded. In addition, a further 6 participants were excluded from this analysis due to missing the resting state recording file (1 subject) or not having enough resting data events to analyze (5 participants) leaving 111 participants included in the final analysis.

In the resting state protocol, participants were instructed to fixate on a central cross, and open or close their eyes when they heard a beep, alternating between 20 second blocks of eyes open and 40 second blocks of eyes closed. The dataset includes a pre-processed and artifact corrected copy of the data, which was used here, with full details of the pre-processing described in (Langer et al., 2017). Briefly, bad electrodes were identified and interpolated, eye artifacts were regressed out of the EEG from EOG electrodes, and a PCA approach was used to remove sparse noise from the data. We further identified flat channels (channels with no data) and interpolated them, and re-referenced data to a common average reference.

For the current analyses, we used the eyes closed resting state data, and extracted the time period of 5 – 35 seconds within the 40 second eyes closed resting segments, excluding the 5 seconds post and prior to eye opening. We used the first block for each participant for analysis. Power spectra were calculated for each channel using Welch’s method, using 2 second windows with 25% overlap.

We then parameterized the calculated power spectra to return estimates of periodic and aperiodic parameters. The model parameterization we used is agnostic to frequency bands, fitting peaks wherever they’re found in the frequency spectrum regardless of canonical band definitions (Haller et al., 2018). We determined that activity was contained in a band if the peak of an oscillation was contained in our aforementioned band definitions. Settings for parameterizing power spectra are as follows: the width for a detected peak was bound between 1 - 8 Hz, with a maximum number of detectable peaks set at 8, a minimum threshold for detecting a peak set at 0.1 au, the threshold for detecting was set at the default value of 2 standard deviations above the noise floor, and spectra were fit in ‘fixed’ mode without a knee.

For all band ratio measures, we calculated Spearman correlations between spectral parameters, including center frequency, power and bandwidth of each oscillation band, as well as the aperiodic exponent, across all channels. We do not report correlations to aperiodic offset, as offset shifts by themselves do not affect ratio measures (see simulation results). In addition, we calculated Spearman correlations between each ratio measure and participants’ ages, and between spectral parameters and age.

## Results

### 3.1 Simulation Results

We started by investigating, in simulation, the extent to which band ratios capture periodic power as typically interpreted, and/or to what extent they are potentially related to other periodic or aperiodic spectral parameters. Measured theta / beta ratios across simulations in which one spectral parameter was changed at a time, are reported in Figure 4. As expected, when examining periodic changes (Figure 4A) the theta / beta ratio is strongly driven by power of theta and beta oscillations. However, ratio measures can also be influenced by the center frequency and bandwidth of the theta and beta peaks. We also replicate previous work showing that the center frequency of the alpha peak can impact measures of theta / beta ratio, (Lansbergen et al., 2011), and extend this to include alpha bandwidth. For aperiodic changes (Figure 4B), we see that the aperiodic exponent has a significant effect on measured ratio values.

**Figure 4.**
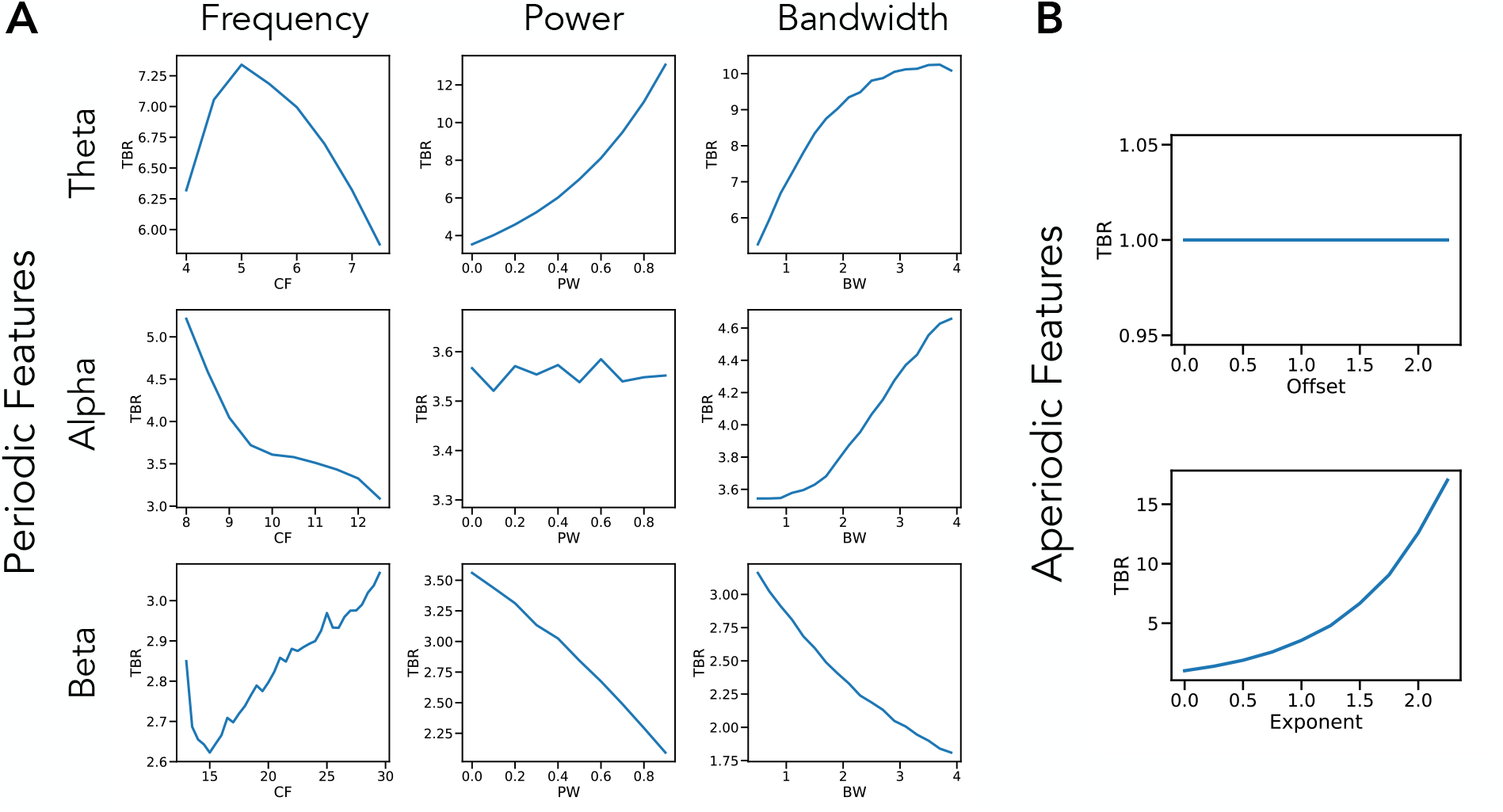
Single Parameter Simulations. Simulations of changes in measured theta / beta ratio as individual parameters are varied, including: **A)** periodic parameters and **B)** aperiodic parameters. Changes in theta center frequency show an increase in theta / beta ratio as the heightened activity is better captured in the canonical band, then decreases as activity leaves the band. Increasing theta power and bandwidth both increase TBR while increasing beta power and bandwidth decreases theta / beta ratio. The center frequency and bandwidth of alpha peaks also influences measured theta / beta ratio, even though alpha is not supposed to be included in the measure. Beta parameters essentially have the inverse effect of changes in theta parameters. Changes in aperiodic exponent also substantially impact measured theta / beta ratio.

Collectively, we see that a wide range of different parameter changes can affect measured ratios. In this case, 8 of the 10 parameters alter theta / beta band ratio, with the only exceptions being the aperiodic offset, which changes power equally between ratio bands, and power in the non-included band, in this case alpha (for the theta / beta ratio). Of note, however, is that the scale of this effects can be quite different, with the power of the included bands and the aperiodic exponent having the biggest impacts. The findings for other band ratio measures are consistent with those for the theta / beta ratio, with full results for them available in the project repository.

We further explored simulations of pairwise combinations of parameter changes, to investigate how ratio measures are affected by concomitant changes in multiple parameters (Figure 5). These simulations include, for example, measured theta / beta band ratios as the aperiodic exponent and theta center frequency both vary, showing an interaction between them (Figure 5A). We can see how changes in aperiodic exponent interact with power changes in the lower (Figure 5B) and higher (Figure 5C) bands. These simulations also demonstrate that both features have an impact on measured ratios, and allow a comparison of scale, showing, for example, that although the influence of low band power and aperiodic exponent is of a similar magnitude, when compared to high band power, the effect of aperiodic exponent changes is relatively much larger. Collectively, through these simulations, we see that changes in different spectral parameters can interact and drive different patterns of differences in measured band ratios. Further simulations of interacting parameters across all other combinations are available in the project repository.

**Figure 5.**
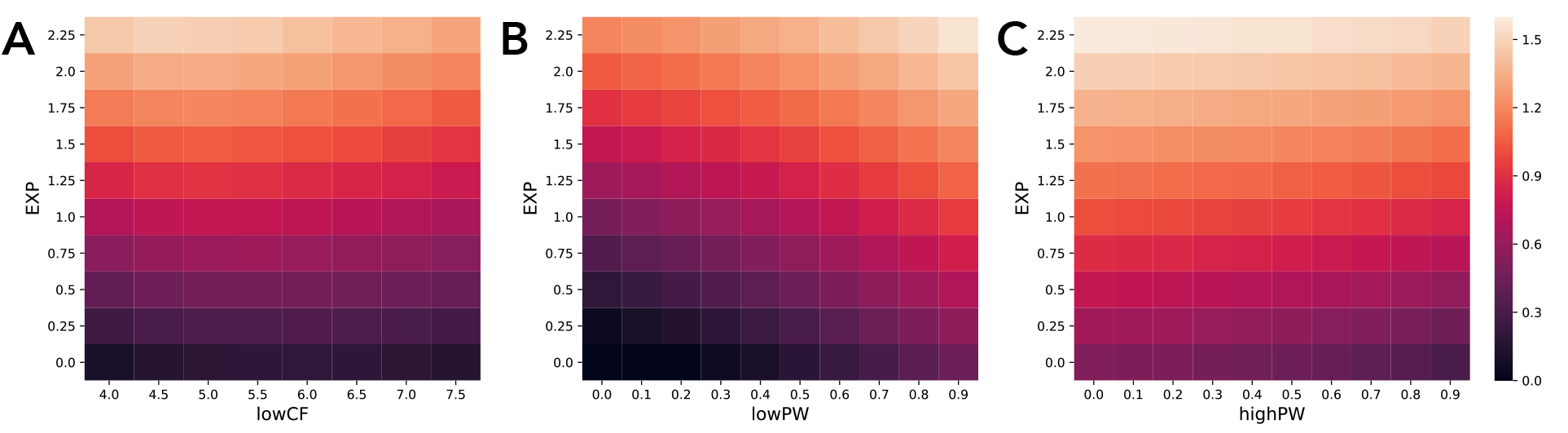
Interacting Parameter Simulations. Measured theta / beta ratio values in simulations as two spectral parameters are varied together. Ratio measures plotted in log10 space due to their skewed distributions. Combinations plotted are aperiodic exponent with low band center frequency **(A)**, as well as with low band power **(B)** and high band power **(C)**. All combinations of varying parameters influence measured band ratio values.

### 3.2 EEG Data Results

We continue our investigation with EEG data recorded during resting state, and compare band ratio measures to parameterized power spectral features. For all correlations here, we report results across all channels. Re-running these analyses with channel groups, using frontal, central, and parietal sub-selections showed qualitatively similar patterns, the results of which are available in the project repository.

For the theta / beta ratio, within periodic spectral parameters we find, as expected, that the strongest relationship is between theta / beta ratio and theta power (*r* = 0.35, *p* < 0.001) with a similarly high correlation with beta power (*r* = −0.29, *p* < 0.01). However, when considering aperiodic parameters, we find a much stronger relationship between theta / beta ratio and aperiodic exponent (*r* = 0.77, *p* < 10^−20^). The full set of spectral parameter correlations is available in Figure 6A.

**Figure 6.**
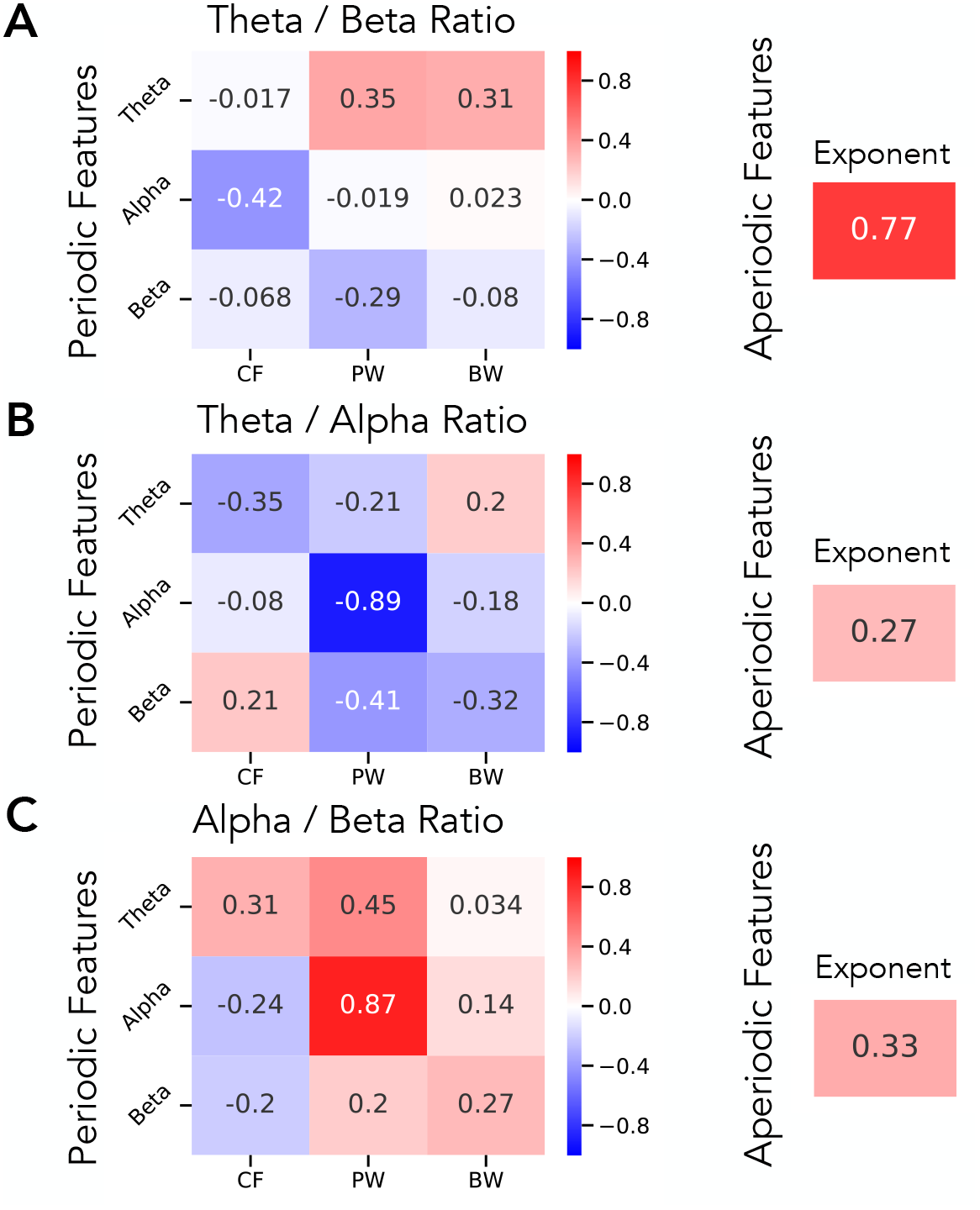
Correlations between Spectral Parameters and Band Ratio Measures in EEG Data. In a large EEG dataset, correlation results are reported for band ratios as compared to the periodic (left) and aperiodic (right) parameters for the **(A)** theta / beta ratio, **(B)** theta / alpha ratio and **(C)** alpha / beta ratio. In **(A)**, these results show that the theta / beta ratio is most strongly correlated with the aperiodic exponent, and less related to power in the theta or beta. In contrast, **(B)** and **(C)** show that any ratio measure that includes an alpha band is most strongly correlated to alpha power, meaning any alpha ratio is mostly reflecting just alpha power.

In contrast, for the theta / alpha ratio, the highest correlation across both periodic and aperiodic spectral parameters was for alpha power (*r* = −0.89, *p* < 10^−35^), with a much lower correlation with aperiodic exponent (*r* = 0.27, *p* < 0.01). This pattern of correlations was also similar for the alpha / beta ratio, with a strong correlation with alpha (*r* = 0.87, *p* < 10^−30^), and a much weaker one with aperiodic exponent (*r* = 0.33, *p* < 0.001). Spectral parameter correlations for the theta / alpha ratio and alpha / beta ratio are available in Figure 6B & 6C respectively.

We also calculated average ratio measures and spectral parameters for each channel, across the group. Topographies of these measures are plotted in Figure 7. Here we can see, for example, that the spatial topography of the theta / beta ratio is most similar to that of the aperiodic exponent, with a strong spatial correlation (*r* = 0.77, *p* < 10^−20^). The topography of alpha / beta ratio is nearly identical to the topography of alpha power (*r* = 0.97, *p* < 10^−70^), with a strong inverse relation between the theta / alpha ratio and alpha power (*r* = −0.92, *p* < 10^−45^).

**Figure 7.**
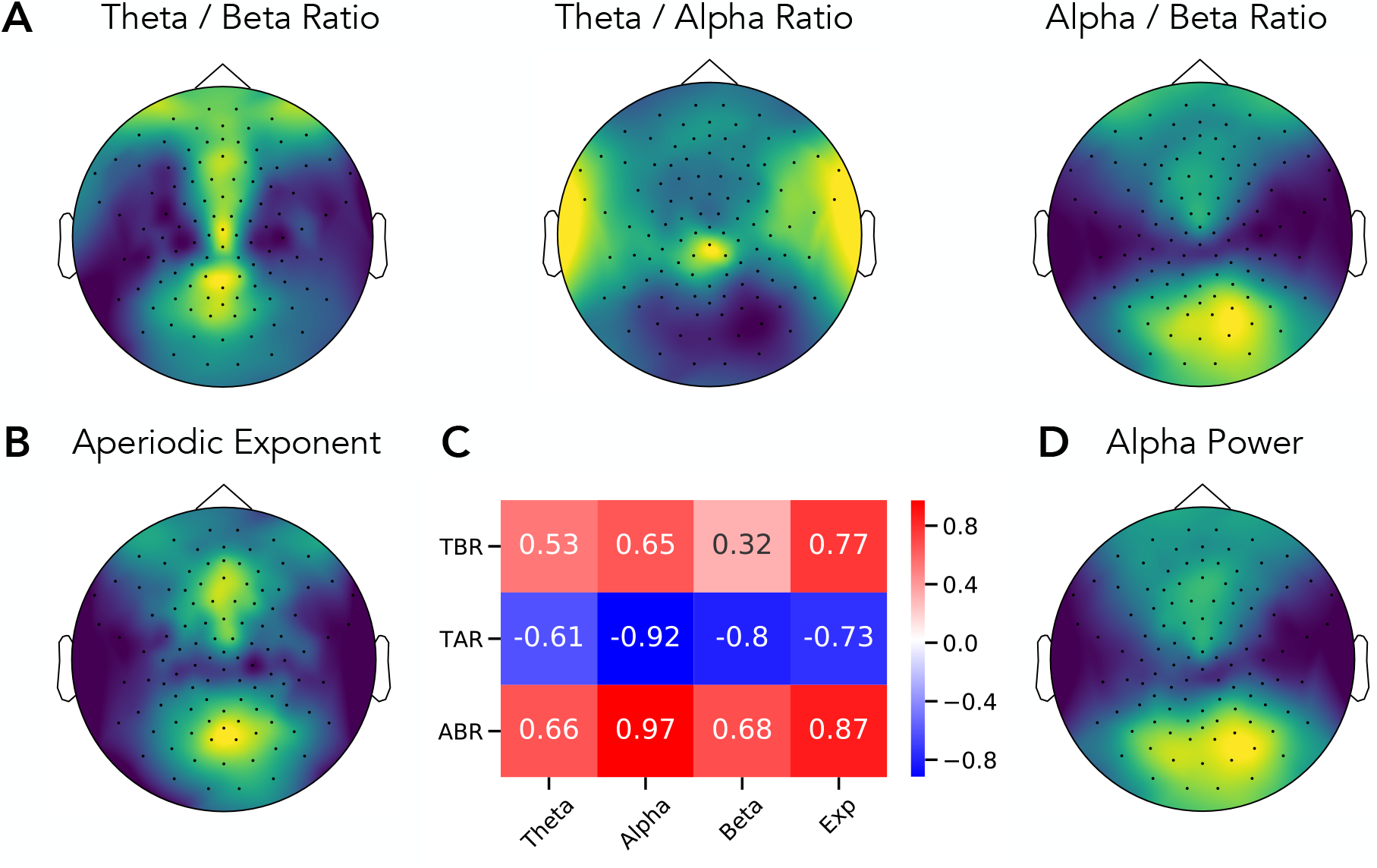
Topographies of Band Ratio Measures and Spectral Parameters. Topographical maps of the **A)** ratios measures, including the theta / beta ratio, theta / alpha ratio and alpha / beta ratio. For comparison, the topography of the aperiodic exponent **(B)** and of alpha power **(D)** are also presented. Each topography is scaled to relative range of the data, with higher values plotted in lighter colors (yellow). **C)** The spatial correlation between topographies of each ratio measure to spectral parameters including power of theta, alpha and beta, and the aperiodic exponent.

We also calculated how each measure correlated with age. The theta / beta ratio was found to be highly correlated with age (*r* = .67, *p* < 10^−15^), with the negative correlation indicating that older adults have higher theta / beta ratios. In comparison, the theta / alpha ratio had a much smaller correlation with age (*r* = −0.37, *p* = 0.0001) and the alpha / beta ratio was not significantly correlated with age (*r* = −0.12, *p* = 0.22). For spectral parameters, the aperiodic exponent was found to be highly correlated with age (*r* = 0.68, *p* < 10^−15^), consistent with previous reports (W. He et al., 2019; Voytek et al., 2015).

## Discussion

### 4.1 Methodological Discussion Points

Through investigations of both simulated and real data, we find that frequency band ratio measures, though typically applied and interpreted as reflecting the relative periodic power of distinct frequency bands, can actually reflect a large number of distinct changes in the underlying data. These band ratio measures therefore capture multiple different changes in periodic and aperiodic properties. Part of this stems from the use of predefined frequency bands of interest, as has been previously reported (Lansbergen et al., 2011; Saad et al., 2018). Here, we replicate and extend this finding, showing how center frequency, and also oscillatory bandwidth, can influence band ratio measures in ways that can be misinterpreted as reflecting power differences. In addition, we show how frequency band ratio measures may commonly capture, at least partially, aperiodic components of electrophysiological data.

Specifically, we used a parameterization model conceiving of the power spectrum as the combination of an aperiodic, 1/f-like spectrum, characterized by an offset and exponent, with overlying periodic ‘peaks’, each characterized by a center frequency, power (over and above the aperiodic background) and bandwidth measure. With this approach, we show many of these parameters can similarly affect band ratio measures in simulation. When applied to real data, we find that different parameters do affect ratio measures, with different patterns for different ratio measures. For example, theta / beta ratio measures mostly reflect aperiodic exponent, whereas theta / alpha and alpha / beta ratios mostly reflect alpha power. In no ratio measures did we find evidence that the measure primarily reflects power within both specified bands.

Given the underdetermined nature of band ratio measures in the face of multiple features of the data that may be changing, we conclude that band ratio measures are not an appropriate measure for characterizing electrophysiological data, at least not in isolation. This is because are uninterpretable in terms of knowing which component(s) of the data they actually reflect. Therefore, we recommend complementary or alternate approaches. These include methods that fully parameterize neural power spectra, specifically measuring periodic and aperiodic components (Haller et al., 2018), which allows for precise quantification of which features of the data vary within and between individuals.

A prior recommendation, that attempts to address center frequency differences (Lansbergen et al., 2011), is that band ratio measures should use individualized frequency bands (Saad et al., 2018). It should be noted that the recommended approach, originally proposed by (Klimesch, 1999), is to use individualized bands based on an alpha band anchor point, whereby theta and beta can be defined as below and above the observed alpha peak. Though this addresses some issues with varying alpha center frequency, it does not specifically establish if there is a defined theta or beta peak, over and above aperiodic power, nor does it identify specific center frequencies should such periodic activity be present. Because this approach also does not separate aperiodic from periodic power, individualized peak detection, especially when anchored to alpha peaks, is insufficient to address the problems highlighted here.

It has previously been reported that ratio measures are stable and have high test-retest reliability within individuals (Angelidis et al., 2016; Monastra et al., 2001; Ohlund, 2000). This is not necessarily in conflict with the finding here that band ratio measures may reflect many distinct features of the data; stable test-retest reliability merely suggests that whichever feature(s) are captured by band ratios within a given subject are themselves stable. However, that band ratios across individuals, and in particular across different populations, may reflect different properties of the data may well help explain why there has been difficulty in reproducing several findings using band ratios. For example, recent failures to replicate band ratio measures include follow ups on previously reported relations with trait anxiety (van Son et al., 2018) or attentional control (van Son et al., 2019). In clinical work, there have been inconsistent findings relating the theta / beta ratio to ADHD (Liechti et al., 2013; Ogrim et al., 2012). It is possible that when investigating varying populations, different features of the data may be driving different observed ratio measures, and this may relate to the significant variance of band ratio measures and their correlates found across studies.

### 4.2 Interpretation Related Discussion Points

The findings cast doubt on the interpretations of prior reports that use band ratio measures and interpret them as primarily reflecting periodic power. Where such studies are reproducible, recontextualization of such findings should consider multiple possible interpretations, including, for example that, a) there is a true change in the power ratio of activity between distinct frequency bands reflecting periodic activity, b) there is a difference in periodic parameters other than power, such as in center frequency and/or bandwidth, c) band ratio measures reflect differences in aperiodic activity, or, d) some combination of the above. Based on data analyzed, the theta / beta ratio is most likely to reflect aperiodic activity, whereas the theta / alpha and alpha / beta ratios are most likely to primarily reflect alpha power. That said, ratio measures could vary across studies in what they reflect, and/or reflect interactions between parameters. Re-evaluations of prior work and/or follow up investigations should seek to re-evaluate such data to investigate which features, in each case, are driving the measured changes in band ratios, and update interpretations accordingly.

In this investigation we replicated the consistently reported finding that band ratio measures vary systematically with age (Angelidis et al., 2016; Bresnahan et al., 1999; Buyck & Wiersema, 2014; Clarke et al., 2001; Gasser et al., 1988; Monastra et al., 2001; Ogrim et al., 2012; Putman et al., 2010), as well as the finding that aperiodic activity also varies systematically with age (Voytek et al., 2015). Since we also find that band ratio measures are highly correlated with aperiodic activity (especially the theta / beta ratio), this is altogether consistent with the idea that the relation of band ratio measures to age is plausibly due to band ratios reflecting aperiodic activity. We note that the dataset used here consists of young participants, and the pattern of findings here is also consistent with recent work showing that the relation of aperiodic activity to age is also apparent in younger participants, and that changes in aperiodic activity across age better explains developmental patterns rather than previous reports of correlated changes across multiple distinct oscillation bands (W. He et al., 2019).

Overall, the EEG data analyzed here suggests that ratio measures, and the theta / beta ratio in particular, often largely reflects aperiodic activity. As well as the relationship of aperiodic activity and band ratios to age, this is also consistent with other reports that previously reported correlates of band ratios have also been found to relate to aperiodic activity. For example, when band ratios are used in sleep scoring, it is typically done with the delta / theta ratio, which we predict likely also captures aperiodic changes, which would be consistent with recent reports that aperiodic activity changes systematically with sleep (Lendner et al., 2019). Collectively, these shared correlates are consistent with suggestion that band ratio measures likely often reflect aperiodic activity.

A key prediction, if ratio measures often reflect aperiodic properties, is that the reported findings will not be specific to the frequency ranges used to measure the ratios, as aperiodic effects should exist across all frequencies. Indeed, correlated change across frequency bands is one of the observations that led to the popularity of band ratio measures (Lubar, 1991). It has also been reported that distinct ratio measures across different frequency bands show similar patterns, for example that both delta / beta and theta / beta ratios relate to cognitive correlates (Schutter & Van Honk, 2005; Tortella-Feliu et al., 2014), both theta / alpha and theta / beta have been reported to relate to ADHD (Barry et al., 2003), and multiple different ratios show similar patterns in investigations of Alzheimer’s disease (Poza et al., 2008). In cases such as these, in which different band ratio measures show approximately similar trends across a wide array of band pairs, a plausible interpretation is that these findings do not reflect correlated changes across multiple distinct frequency bands, but rather that they are all capturing frequency-agnostic aperiodic shifts.

In neurofeedback designs, where band ratios are a target for manipulation rather than a descriptive measure, findings are also consistent with the possibility that targeting ratios at least partially manipulates aperiodic properties, rather than targeting oscillation bands specifically. For example, a recent report showed that targeting beta in a feedback design also induces changes in the alpha band (Jurewicz et al., 2018), which challenges the possibility of targeting different bands independently. Where investigations probe the specificity of neurofeedback protocols, non-specific effects have been reported, such as an effect on beta from a theta / alpha protocol (Egner et al., 2004), and changes in alpha when using a theta / beta protocol (Bazanova et al., 2018; Limin Yang et al., 2015), all of which is consistent with ratios reflecting aperiodic activity.

If a considerable proportion of the variance of band ratios measures is due to aperiodic properties, and not well described or interpreted as band specific changes, then it becomes an open question to ask what the physiological interpretation should be, and therefore how these findings should be interpreted. One hypothesis is that the aperiodic properties of neural time series may relate the relative balance of excitatory and inhibitory activity (Gao et al., 2017). Though further work is required to explore this hypothesis and how it relates to measurements done with band ratios, this does suggest a potential link between what has been measured in band ratios, as a correlate of various cognitive markers and disease states, and potential interpretations related to excitation and inhibition. A more general review of aperiodic properties in neural data, sometimes referred to ‘scale-free’ activity, is available in (B. J. He, 2014).

Particular attention should be paid to ratio measures applied in clinical applications, in which the pursuit of biomarkers based on faulty measures could hinder, rather than ameliorate, clinical practice. For example, the findings here on ratio measures are consistent with the practice advisory that using theta / beta ratio measures in the context of ADHD is not an appropriate measure (Gloss et al., 2016). Rather, the prediction based on these results for ADHD would be that the oft reported theta / beta correlate is likely a reflection of differences in aperiodic activity. In other work, we have found exactly this: that aperiodic properties are correlated with theta / beta measures in a population with ADHD, and that the aperiodic measures themselves better relate not only to disease state but also to medication status (Robertson et al., 2019). For other clinical disorders that have been investigated with band ratio measures, such as Alzheimer’s disease (Cassani et al., 2018), or psychotic disorders (Howells et al., 2018) we recommend that investigations should follow up on which underlying features best explain changes in ratio measures, and update interpretations and future work on biomarkers accordingly.

A notable exception, as we found in analyzed EEG data, to ratio measures reflecting aperiodic shifts is in cases in which ratio measures include the alpha band. When the alpha band is included in the ratio, band ratio measures tend to primarily reflect alpha power. This is likely due to the prominence of the alpha band, where alpha is typically present across participants, has very high power, and is dynamic. Thus, it is logical that ratio measures that include the alpha band largely reflect alpha dynamics, as we observed here. This effect may also be exaggerated in our analysis, as we are analyzing eyes closed data, in which alpha power is most prominent, though the pattern of results is consistent when re-run on eyes open data. Investigations in which ratio measures such as delta / alpha or theta / alpha are used should investigate to what extent the dominant effect they are capturing is alpha dynamics. Overall we recommend that reports from studies using band ratios including alpha should consider if the findings are likely to be largely explained by alpha dynamics.

## Conclusion

Frequency band ratio measures are a common analysis approach applied to neural field data, including EEG, MEG, ECoG and LFP. Band ratio approaches have been applied across many domains, including basic research investigating executive functions, learning and memory, and sleep; in clinical investigations including investigating ADHD and dementia; and in applied work leveraging them for neurofeedback applications. Though typically interpreted as a normalized measure reflecting the relative power of distinct periodic components, here we show that band ratio measures can reflect not only multiple features of periodic neural activity, including the center frequency, power and bandwidth of periodic components, but can also be driven by variations in aperiodic activity. This is demonstrated in simulation, and also in empirical work applied to a large EEG dataset in which we show how multiple spectral features relate to measured band ratios, making them an imprecise metric. For example, the most dominant contributor to the theta / beta ratio is the aperiodic exponent, whereas the theta / alpha and alpha / beta ratio predominantly reflect alpha power. Overall, band ratio measures are found to be underdetermined, and so across participants, recording modalities, species, and contexts may reflect different components of the signal. This makes comparisons with band ratio measures difficult, if not impossible, and questions their typical interpretations as reflecting periodic activity. As an alternative, we recommend that parameterization of neural power spectra is able to better capture which components of neural signals vary and relate to features of interest, without conflating changes in periodic and aperiodic activity, as band ratio measures do.

## Acknowledgements

We would like to thank members of the Voytek Lab for insightful comments and suggestions throughout this project. We would also like to express gratitude to the many people involved in generating the open-access datasets and developing the open-source tools that made this project possible.

## Abbreviations

EEG: electroencephalography
MEG: magnetoencephalography
ECoG: electrocorticography
LFP: local field potential
TBR: theta / beta ratio
TAR: theta / alpha ratio
ABR: alpha / beta ratio
CF: center frequency
PW: power
BW: bandwidth
EXP: aperiodic exponent
ADHD: attention-deficit hyperactivity disorder

## Supplementary Materials

**Supplemental Table 1.** Simulation Parameters for Periodic Components

|  |  | **Theta** | **Alpha** | **Beta** |
| --- | --- | --- | --- | --- |
| **CF** | Default | 6 | 10 | 21.5 |
|  | Range | 4 - 8 | 8 - 13 | 13 - 30 |
|  | Increment | 0.25 | 0.25 | 1 |
| **PW** | Default | 0.5 | 0.5 | 0.5 |
|  | Range | 0 - 1.0 | 0 - 1.0 | 0 - 1.0 |
|  | Increment | 0.1 | 0.1 | 0.1 |
| **BW** | Default | 0.1 | 0.1 | 0.1 |
|  | Range | 0.2 - 0.4 | 0.2 - 0.4 | 0.2 - 0.4 |
|  | Increment | 0.2 | 0.2 | 0.2 |

**Supplemental Table 2.** Simulation Parameters for Aperiodic Components

|  | **Default** | **Range** | **Increment** |
| --- | --- | --- | --- |
| **Offset** | 0 | 0 - 2.5 | 0.25 |
| **Exponent** | 1 | 0 - 3 | 0.2 |

## References

Angelidis, A., Hagenaars, M., van Son, D., van der Does, W., & Putman, P. (2018). Do not look away! Spontaneous frontal EEG theta/beta ratio as a marker for cognitive control over attention to mild and high threat. Biological Psychology, 135, 8–17. https://doi.org/10.1016/j.biopsycho.2018.03.002

Angelidis, A., van der Does, W., Schakel, L., & Putman, P. (2016). Frontal EEG theta/beta ratio as an electrophysiological marker for attentional control and its test-retest reliability. Biological Psychology, 121, 49–52. https://doi.org/10.1016/j.biopsycho.2016.09.008

Arns, M., Conners, C. K., & Kraemer, H. C. (2013). A Decade of EEG Theta/Beta Ratio Research in ADHD: A Meta-Analysis. Journal of Attention Disorders, 17(5), 374–383. https://doi.org/10.1177/1087054712460087

Arns, M., Heinrich, H., & Strehl, U. (2014). Evaluation of neurofeedback in ADHD: The long and winding road. Biological Psychology, 95, 108–115. https://doi.org/10.1016/j.biopsycho.2013.11.013

Barry, R. J., Clarke, A. R., & Johnstone, S. J. (2003). A review of electrophysiology in attention-deficit/hyperactivity disorder: I. Qualitative and quantitative electroencephalography. Clinical Neurophysiology, 114(2), 171–183. https://doi.org/10.1016/S1388-2457(02)00362-0

Bazanova, O. M., Auer, T., & Sapina, E. A. (2018). On the Efficiency of Individualized Theta/Beta Ratio Neurofeedback Combined with Forehead EMG Training in ADHD Children. Frontiers in Human Neuroscience, 12. https://doi.org/10.3389/fnhum.2018.00003

Bennys, K., Rondouin, G., Vergnes, C., & Touchon, J. (2001). Diagnostic value of quantitative EEG in Alzheimer’s disease. Neurophysiologie Clinique, 31(3), 153–160. https://doi.org/10.1016/S0987-7053(01)00254-4

Bresnahan, S. M., Anderson, J. W., & Barry, R. J. (1999). Age-related changes in quantitative EEG in attention-deficit / hyperactivity disorder. Biological Psychiatry, 46(12), 1690–1697. https://doi.org/10.1016/S0006-3223(99)00042-6

Buyck, I., & Wiersema, J. R. (2014). State-related electroencephalographic deviances in attention deficit hyperactivity disorder. Research in Developmental Disabilities, 35(12), 3217–3225. https://doi.org/10.1016/j.ridd.2014.08.003

Buzsáki, G., Logothetis, N., & Singer, W. (2013). Scaling Brain Size, Keeping Timing: Evolutionary Preservation of Brain Rhythms. Neuron, 80(3), 751–764. https://doi.org/10.1016/j.neuron.2013.10.002

Cassani, R., Estarellas, M., San-Martin, R., Fraga, F. J., & Falk, T. H. (2018). Systematic Review on Resting-State EEG for Alzheimer’s Disease Diagnosis and Progression Assessment. Disease Markers, 2018, 1–26. https://doi.org/10.1155/2018/5174815

Clarke, A. R., Barry, R. J., Dupuy, F. E., McCarthy, R., Selikowitz, M., & Johnstone, S. J. (2013). Excess beta activity in the EEG of children with attention-deficit/hyperactivity disorder: A disorder of arousal? International Journal of Psychophysiology, 89(3), 314–319. https://doi.org/10.1016/j.ijpsycho.2013.04.009

Clarke, A. R., Barry, R. J., McCarthy, R., & Selikowitz, M. (2001). Age and sex effects in the EEG: Development of the normal child. Clinical Neurophysiology, 112(5), 806–814. https://doi.org/10.1016/S1388-2457(01)00488-6

Costa-Miserachs, D., Portell-Cortés, I., Torras-Garcia, M., & Morgado-Bernal, I. (2003). Automated sleep staging in rat with a standard spreadsheet. Journal of Neuroscience Methods, 130(1), 93–101. https://doi.org/10.1016/S0165-0270(03)00229-2

Daniel, R. S. (1964). Electroencephalographic Correlogram Ratios and Their Stability. Science, 145(3633), 721–723. https://doi.org/10.1126/science.145.3633.721

Donoghue, T. (2019). LISC: A Python Package for Scientific Literature Collection and Analysis. Journal of Open Source Software, 4(41), 1674. https://doi.org/10.21105/joss.01674

Egner, T., Strawson, E., & Gruzelier, J. H. (2002). EEG Signature and Phenomenology of Alpha/theta Neurofeedback Training Versus Mock Feedback. Applied Psychophysiology and Biofeedback, 27(4), 261–270. https://doi.org/10.1023/A:1021063416558

Egner, T., Zech, T. F., & Gruzelier, J. H. (2004). The effects of neurofeedback training on the spectral topography of the electroencephalogram. Clinical Neurophysiology, 115(11), 2452–2460. https://doi.org/10.1016/j.clinph.2004.05.033

Gao, R., Peterson, E. J., & Voytek, B. (2017). Inferring synaptic excitation/inhibition balance from field potentials. NeuroImage, 158, 70–78. https://doi.org/10.1016/j.neuroimage.2017.06.078

Gasser, T., Verleger, R., Bächer, P., & Sroka, L. (1988). Development of the EEG of school-age children and adolescents. I. Analysis of band power. Electroencephalography and Clinical Neurophysiology, 69(2), 91–99. https://doi.org/10.1016/0013-4694(88)90204-0

Geraedts, V. J., Marinus, J., Gouw, A. A., Mosch, A., Stam, C. J., van Hilten, J. J., Contarino, M. F., & Tannemaat, M. R. (2018). Quantitative EEG reflects non-dopaminergic disease severity in Parkinson’s disease. Clinical Neurophysiology, 129(8), 1748–1755. https://doi.org/10.1016/j.clinph.2018.04.752

Gloss, D., Varma, J. K., Pringsheim, T., & Nuwer, M. R. (2016). Practice advisory: The utility of EEG theta/beta power ratio in ADHD diagnosis: Report of the Guideline Development, Dissemination, and Implementation Subcommittee of the American Academy of Neurology. Neurology, 87(22), 2375–2379. https://doi.org/10.1212/WNL.0000000000003265

Gordon, S., Todder, D., Deutsch, I., Garbi, D., Getter, N., & Meiran, N. (2018). Are resting state spectral power measures related to executive functions in healthy young adults? Neuropsychologia, 108, 61–72. https://doi.org/10.1016/j.neuropsychologia.2017.10.031

Gramfort, A., Luessi, M., Larson, E., Engemann, D. A., Strohmeier, D., Brodbeck, C., Parkkonen, L., & Hämäläinen, M. S. (2014). MNE software for processing MEG and EEG data. NeuroImage, 86, 446–460. https://doi.org/10.1016/j.neuroimage.2013.10.027

Haller, M., Donoghue, T., Peterson, E., Varma, P., Sebastian, P., Gao, R., Noto, T., Knight, R. T., Shestyuk, A., & Voytek, B. (2018). Parameterizing neural power spectra. BioRxiv. https://doi.org/10.1101/299859

He, B. J. (2014). Scale-free brain activity: Past, present, and future. Trends in Cognitive Sciences, 18(9), 480–487. https://doi.org/10.1016/j.tics.2014.04.003

He, W., Donoghue, T., Sowman, P. F., Seymour, R. A., Brock, J., Crain, S., Voytek, B., & Hillebrand, A. (2019). Co-Increasing Neuronal Noise and Beta Power in the Developing Brain. BioRxiv, 49.

Howells, F. M., Temmingh, H. S., Hsieh, J. H., van Dijen, A. V., Baldwin, D. S., & Stein, D. J. (2018). Electroencephalographic delta/alpha frequency activity differentiates psychotic disorders: A study of schizophrenia, bipolar disorder and methamphetamine-induced psychotic disorder. Translational Psychiatry, 8(1). https://doi.org/10.1038/s41398-018-0105-y

Jurewicz, K., Paluch, K., Kublik, E., Rogala, J., Mikicin, M., & Wróbel, A. (2018). EEG-neurofeedback training of beta band (12–22 Hz) affects alpha and beta frequencies – A controlled study of a healthy population. Neuropsychologia, 108, 13–24. https://doi.org/10.1016/j.neuropsychologia.2017.11.021

Keune, P. M., Hansen, S., Weber, E., Zapf, F., Habich, J., Muenssinger, J., Wolf, S., Schönenberg, M., & Oschmann, P. (2017). Exploring resting-state EEG brain oscillatory activity in relation to cognitive functioning in multiple sclerosis. Clinical Neurophysiology, 128(9), 1746–1754. https://doi.org/10.1016/j.clinph.2017.06.253

Kim, J., Goldsberry, M. E., Harmon, T. C., & Freeman, J. H. (2016). Developmental Changes in Hippocampal CA1 Single Neuron Firing and Theta Activity during Associative Learning. PLOS ONE, 11(10), e0164781. https://doi.org/10.1371/journal.pone.0164781

Klimesch, W. (1999). EEG alpha and theta oscillations reflect cognitive and memory performance: A review and analysis. Brain Research Reviews, 29(2), 169–195. https://doi.org/10.1016/S0165-0173(98)00056-3

Krakovská, A., & Mezeiová, K. (2011). Automatic sleep scoring: A search for an optimal combination of measures. Artificial Intelligence in Medicine, 53(1), 25–33. https://doi.org/10.1016/j.artmed.2011.06.004

Langer, N., Ho, E. J., Alexander, L. M., Xu, H. Y., Jozanovic, R. K., Henin, S., Petroni, A., Cohen, S., Marcelle, E. T., Parra, L. C., Milham, M. P., & Kelly, S. P. (2017). A resource for assessing information processing in the developing brain using EEG and eye tracking. Scientific Data, 4, 170040. https://doi.org/10.1038/sdata.2017.40

Lansbergen, M. M., Arns, M., van Dongen-Boomsma, M., Spronk, D., & Buitelaar, J. K. (2011). The increase in theta/beta ratio on resting-state EEG in boys with attention-deficit/hyperactivity disorder is mediated by slow alpha peak frequency. Progress in Neuro-Psychopharmacology and Biological Psychiatry, 35(1), 47–52. https://doi.org/10.1016/j.pnpbp.2010.08.004

Lendner, J. D., Helfrich, R. F., Mander, B. A., Romundstad, L., Lin, J. J., Walker, M. P., Larsson, P. G., & Knight, R. T. (2019). An Electrophysiological Marker of Arousal Level in Humans. BioRxiv. https://doi.org/10.1101/625210

Liechti, M. D., Valko, L., Müller, U. C., Döhnert, M., Drechsler, R., Steinhausen, H.-C., & Brandeis, D. (2013). Diagnostic Value of Resting Electroencephalogram in Attention-Deficit/Hyperactivity Disorder Across the Lifespan. Brain Topography, 26(1), 135–151. https://doi.org/10.1007/s10548-012-0258-6

Limin Yang, Wenya Nan, Xiaoting Qu, Feng Wan, Pui-In Mak, Peng Un Mak, Vai, M. I., Yong Hu, & Rosa, A. (2015). Beta/theta ratio neurofeedback training effects on the spectral topography of EEG. 37th Annual International Conference of the IEEE Engineering in Medicine and Biology Society (EMBC), 4741–4744. https://doi.org/10.1109/EMBC.2015.7319453

Long, C. W., Shah, N. K., Loughlin, C., Spydell, J., & Bedford, R. F. (1989). A Comparison of EEG Determinants of Near-Awakening from Isoflurane and Fentanyl Anesthesia: Spectral Edge, Median Power Frequency, and δ Ratio. Anesthesia & Analgesia, 69(2), 169–173. https://doi.org/10.1213/00000539-198908000-00005

Loo, S. K., & Makeig, S. (2012). Clinical Utility of EEG in Attention-Deficit/Hyperactivity Disorder: A Research Update. Neurotherapeutics, 9(3), 569–587. https://doi.org/10.1007/s13311-012-0131-z

Lubar, J. F. (1991). Discourse on the development of EEG diagnostics and biofeedback for attention-deficit/hyperactivity disorders. Biofeedback and Self-Regulation, 16(3), 201–225. https://doi.org/10.1007/BF01000016

Matoušek, M. (1968). Frequency Analysis in Routine Electroencephalography. Electroencephalography and Clinical Neurophysiology, 24(4), 365–373. https://doi.org/10.1016/0013-4694(68)90197-1

Matoušek, M., & Petersén, I. (1973). Automatic evaluation of EEG background activity by means of age-dependent EEG quotients. Electroencephalography and Clinical Neurophysiology, 35(6), 603–612. https://doi.org/10.1016/0013-4694(73)90213-7

Matoušek, M., & Petersén, I. (1983). A method for assessing alertness fluctuations from EEG spectra. Electroencephalography and Clinical Neurophysiology, 55(1), 108–113. https://doi.org/10.1016/0013-4694(83)90154-2

Monastra, V. J., Lubar, J. F., & Linden, M. (2001). The development of a quantitative electroencephalographic scanning process for attention deficit-hyperactivity disorder: Reliability and validity studies. Neuropsychology, 15(1), 136–144. https://doi.org/10.1037//0894-4105.15.1.136

Moretti, D. V., Fracassi, C., Pievani, M., Geroldi, C., Binetti, G., Zanetti, O., Sosta, K., Rossini, P. M., & Frisoni, G. B. (2009). Increase of theta/gamma ratio is associated with memory impairment. Clinical Neurophysiology, 120(2), 295–303. https://doi.org/10.1016/j.clinph.2008.11.012

Moretti, D. V., Paternicò, D., Binetti, G., Zanetti, O., & Frisoni, G. B. (2013). EEG upper/low alpha frequency power ratio relates to temporo-parietal brain atrophy and memory performances in mild cognitive impairment. Frontiers in Aging Neuroscience, 5. https://doi.org/10.3389/fnagi.2013.00063

Nokia, M. S., Penttonen, M., Korhonen, T., & Wikgren, J. (2008). Hippocampal theta (3–8Hz) activity during classical eyeblink conditioning in rabbits. Neurobiology of Learning and Memory, 90(1), 62–70. https://doi.org/10.1016/j.nlm.2008.01.005

Ogrim, G., Kropotov, J., & Hestad, K. (2012). The quantitative EEG theta/beta ratio in attention deficit/hyperactivity disorder and normal controls: Sensitivity, specificity, and behavioral correlates. Psychiatry Research, 198(3), 482–488. https://doi.org/10.1016/j.psychres.2011.12.041

Ohlund, B. (2000). An Investigation of the Reliability and Validity of Theta/Beta Ratio Measurement [PhD Thesis]. Arizona State University.

Penttilä, M., Partanen, J. V., Soininen, H., & Riekkinen, P. J. (1985). Quantitative analysis of occipital EEG in different stages of Alzheimer’s disease. Electroencephalography and Clinical Neurophysiology, 60(1), 1–6. https://doi.org/10.1016/0013-4694(85)90942-3

Pfurtscheller, G., Schwarz, G., & List, W. (1986). Long-lasting EEG reactions in comatose patients after repetitive stimulation. Electroencephalography and Clinical Neurophysiology, 64, 402–410. https://doi.org/10.1016/0013-4694(86)90073-8

Podvalny, E., Noy, N., Harel, M., Bickel, S., Chechik, G., Schroeder, C. E., Mehta, A. D., Tsodyks, M., & Malach, R. (2015). A unifying principle underlying the extracellular field potential spectral responses in the human cortex. Journal of Neurophysiology, 114(1), 505–519. https://doi.org/10.1152/jn.00943.2014

Poza, J., Hornero, R., Abásolo, D., Fernández, A., & Mayo, A. (2008). Evaluation of spectral ratio measures from spontaneous MEG recordings in patients with Alzheimer’s disease. Computer Methods and Programs in Biomedicine, 90(2), 137–147. https://doi.org/10.1016/j.cmpb.2007.12.004

Putman, P., van Peer, J., Maimari, I., & van der Werff, S. (2010). EEG theta/beta ratio in relation to fear-modulated response-inhibition, attentional control, and affective traits. Biological Psychology, 83(2), 73–78. https://doi.org/10.1016/j.biopsycho.2009.10.008

Raymond, J., Varney, C., Parkinson, L. A., & Gruzelier, J. H. (2005). The effects of alpha/theta neurofeedback on personality and mood. Cognitive Brain Research, 23(2–3), 287–292. https://doi.org/10.1016/j.cogbrainres.2004.10.023

Reed, C. M., Birch, K. G., Kamiński, J., Sullivan, S., Chung, J. M., Mamelak, A. N., & Rutishauser, U. (2017). Automatic detection of periods of slow wave sleep based on intracranial depth electrode recordings. Journal of Neuroscience Methods, 282, 1–8. https://doi.org/10.1016/j.jneumeth.2017.02.009

Robertson, M. M., Furlong, S., Voytek, B., Donoghue, T., Boettiger, C. A., & Sheridan, M. A. (2019). EEG Power Spectral Slope differs by ADHD status and stimulant medication exposure in early childhood. Journal of Neurophysiology. https://doi.org/10.1152/jn.00388.2019

Saad, J. F., Kohn, M. R., Clarke, S., Lagopoulos, J., & Hermens, D. F. (2018). Is the Theta/Beta EEG Marker for ADHD Inherently Flawed? Journal of Attention Disorders, 22(9), 815–826. https://doi.org/10.1177/1087054715578270

Schutter, D. J. L. G., & Van Honk, J. (2005). Electrophysiological ratio markers for the balance between reward and punishment. Cognitive Brain Research, 24(3), 685–690. https://doi.org/10.1016/j.cogbrainres.2005.04.002

Sheorajpanday, R. V. A., Nagels, G., Weeren, A. J. T. M., van Putten, M. J. A. M., & De Deyn, P. P. (2009). Reproducibility and clinical relevance of quantitative EEG parameters in cerebral ischemia: A basic approach. Clinical Neurophysiology, 120(5), 845–855. https://doi.org/10.1016/j.clinph.2009.02.171

Snyder, S. M., & Hall, J. R. (2006). A Meta-analysis of Quantitative EEG Power Associated With Attention-Deficit Hyperactivity Disorder: Journal of Clinical Neurophysiology, 23(5), 441–456. https://doi.org/10.1097/01.wnp.0000221363.12503.78

Snyder, S. M., Rugino, T. A., Hornig, M., & Stein, M. A. (2015). Integration of an EEG biomarker with a clinician’s ADHD evaluation. Brain and Behavior, 5(4), e00330. https://doi.org/10.1002/brb3.330

Studer, P., Kratz, O., Gevensleben, H., Rothenberger, A., Moll, G. H., Hautzinger, M., & Heinrich, H. (2014). Slow cortical potential and theta/beta neurofeedback training in adults: Effects on attentional processes and motor system excitability. Frontiers in Human Neuroscience, 8. https://doi.org/10.3389/fnhum.2014.00555

Tortella-Feliu, M., Morillas-Romero, A., Balle, M., Llabrés, J., Bornas, X., & Putman, P. (2014). Spontaneous EEG activity and spontaneous emotion regulation. International Journal of Psychophysiology, 94(3), 365–372. https://doi.org/10.1016/j.ijpsycho.2014.09.003

Trammell, J. P., MacRae, P. G., Davis, G., Bergstedt, D., & Anderson, A. E. (2017). The Relationship of Cognitive Performance and the Theta-Alpha Power Ratio Is Age-Dependent: An EEG Study of Short Term Memory and Reasoning during Task and Resting-State in Healthy Young and Old Adults. Frontiers in Aging Neuroscience, 9. https://doi.org/10.3389/fnagi.2017.00364

van Luijtelaar, E., & Coenen, A. (1984). An EEG averaging technique for automated sleep-wake stage identification in the rat. Physiology & Behavior, 33(5), 837–841. https://doi.org/10.1016/0031-9384(84)90056-8

van Son, D., De Blasio, F. M., Fogarty, J. S., Angelidis, A., Barry, R. J., & Putman, P. (2019). Frontal EEG theta/beta ratio during mind wandering episodes. Biological Psychology, 140, 19–27. https://doi.org/10.1016/j.biopsycho.2018.11.003

van Son, D., Schalbroeck, R., Angelidis, A., van der Wee, N. J. A., van der Does, W., & Putman, P. (2018). Acute effects of caffeine on threat-selective attention: Moderation by anxiety and EEG theta/beta ratio. Biological Psychology, 136, 100–110. https://doi.org/10.1016/j.biopsycho.2018.05.006

Vernon, D., Egner, T., Cooper, N., Compton, T., Neilands, C., Sheri, A., & Gruzelier, J. (2003). The effect of training distinct neurofeedback protocols on aspects of cognitive performance. International Journal of Psychophysiology, 47(1), 75–85. https://doi.org/10.1016/S0167-8760(02)00091-0

Voytek, B., Kramer, M. A., Case, J., Lepage, K. Q., Tempesta, Z. R., Knight, R. T., & Gazzaley, A. (2015). Age-Related Changes in 1/f Neural Electrophysiological Noise. Journal of Neuroscience, 35(38), 13257–13265. https://doi.org/10.1523/JNEUROSCI.2332-14.2015

Wang, Y., Sokhadze, E. M., El-Baz, A. S., Li, X., Sears, L., Casanova, M. F., & Tasman, A. (2016). Relative Power of Specific EEG Bands and Their Ratios during Neurofeedback Training in Children with Autism Spectrum Disorder. Frontiers in Human Neuroscience, 9. https://doi.org/10.3389/fnhum.2015.00723

